# Data-Independent Acquisition (DIA)-Based Label-Free Redox Proteomics (DIALRP) Reveals Novel Oxidative Stress Responsive Translation Factors

**DOI:** 10.1101/2024.12.16.626735

**Authors:** Daiki Kobayashi, Tomoyo Takami, Masaki Matsumoto

## Abstract

Oxidative stress is a key factor in numerous physiological and pathological processes, including aging, cancer, and neurodegenerative diseases. Protein cysteine residues are particularly susceptible to oxidative stress-induced modifications that can alter their structure and function, thereby affecting intracellular signaling pathways. In this study, we developed a data-independent acquisition mass spectrometry (DIA-MS)-based label-free redox proteomics method, termed DIALRP, to comprehensively analyse cysteine oxidative modifications in the prostate cancer cell line DU145 under oxidative stress induced by menadione (MND). Through these analyses, we identified translation-related factors with significantly elevated cysteine oxidation upon MND treatment and evaluated their functional relevance. Notably, our data demonstrated that the inhibition of EIF2, EIF6, and EEF2 complex formation due to oxidative stress occurs during the cellular response to translational inhibition. These insights reveal a previously unrecognized mechanism of translation regulation under oxidative stress and provide a valuable framework for future studies on redox-mediated cellular processes.

## Introduction

Oxidative stress is a hallmark of various physiological and pathological processes, including aging, cancer, and neurodegenerative diseases(Valko *et al*, 2007; Reuter *et al*, 2010; Lin & Beal, 2006). Reactive oxygen species (ROS), the major mediators of oxidative stress, when produced in excess, disrupt cellular homeostasis by oxidizing lipids, nucleic acids, and proteins, leading to functional changes and cellular damage. In particular, proteins can undergo oxidative modifications, resulting in conformational changes, altered enzyme activity, and perturbation of cellular signaling pathways(Lennicke & Cochemé, 2021). One of the major oxidative modifications of proteins occurs at cysteine residues, resulting in reversible and irreversible oxidative post-translational modifications of cysteine(Paulsen & Carroll, 2013). Reversible modifications alter the structure and function of proteins(Rhee, 2006; D’Autréaux & Toledano, 2007; Poole & Nelson, 2008; Lennicke & Cochemé, 2021) and several studies have demonstrated their functional roles in signal transduction(Heppner *et al*, 2018), metabolic adaptation(Van Der Reest *et al*, 2018), and developmental hematopoiesis(Pimkova *et al*, 2022). However, modifications of specific cysteines can have diverse consequences depending on the target and type; hence, the effects of ROS-induced protein modifications on cellular signaling are not yet fully understood.

Redox proteomics has emerged as a powerful approach to systematically identify and quantify protein cysteine oxidative modifications, providing insights into cellular responses to oxidative stress. Particularly in cancer biology, it could be useful for a comprehensive analysis of the oxidative stress response. Cancer cells are highly susceptible to oxidative stress, which significantly affects their survival, proliferation, and metastatic potential(Moloney & Cotter, 2018; Wang *et al*, 2021). Prostate cancer is one type of cancer in which the relationship between malignant transformation and ROS has been reported(Kumar *et al*, 2008). In particular, DU145 is an Nrf2-dependent cell line that adapts to stress responses and is used as a cancer cell model to analyse oxidative stress responses(Jayakumar *et al*, 2014; Tossetta *et al*, 2023).

Redox proteomics is often performed using isobaric tag reagents, such as tandem mass tags (TMT). To enhance comprehensiveness, cysteine-containing peptides were subjected to click reaction(Desai *et al*, 2022), anti-iodoTMT antibody-(Pimkova *et al*, 2022), and CPT tag-(Xiao *et al*, 2020)based enrichment methods to increase the number of identified peptides. A cysteine-labeling method using stable isotope-labeled iodoacetamide has also been reported(Van Der Reest *et al*, 2018). This method does not enrich cysteine peptides, but instead improves peptide identification by peptide fractionation. These methods require sophisticated protocols, considerable sample volumes for the enrichment of cysteine-containing peptides, and expensive stable isotope-labeling reagents. To solve these problems, an approach using label-free and data-independent acquisition (DIA)/SWATH-based quantification without peptide enrichment has been reported to profile the cysteine redox status in the supernatants of H_2_O_2_-treated cultured cells(Anjo *et al*, 2019). Thus, redox proteomic analysis methods have been improved and applied to analyze the biological processes related to oxidative stress.

In this study, we developed a novel method for redox proteomic analysis using data-independent acquisition mass spectrometry (DIA-MS) and measured global cysteine redox changes in DU145 cells under oxidative stress conditions caused by menadione (MND). MND is a widely used reagent that generates ROS during oxidative metabolism(Thor *et al*, 1982), allowing researchers to investigate cellular responses to intrinsic oxidative stress in a controlled manner(Chiou & Tzeng, 2000). Consequently, we identified several translation-related factors undergoing oxidative modification following MND treatment, providing new insights into the molecular mechanisms regulating protein synthesis in response to oxidative stress.

## Results and Discussion

### Overview of DIALRP and its application to the analysis of menadione response in DU145 cells

Mass spectrometry-based methods for redox proteomic analysis rely primarily on stable isotopes(Knoke & Leichert, 2023). In the present study, we developed a label-free redox proteomic analysis method based on data-independent acquisition mass spectrometry (DIA-MS) (Fig.1A). Redox proteomics methods require the almost complete removal of excess cysteine-modifying reagents. As previously reported(Desai *et al*, 2022), the single-pot solid-phase enhanced sample pretreatment (SP3) method was also applied to this method(Hughes *et al*, 2019). Each cell lysate was divided into two portions: one where the reduced cysteines were modified with N-ethylmaleimide (NEM), and the other was left unmodified. Proteins were captured using magnetic beads, washed to remove residual reagents, and digested with trypsin. After digestion, both samples were treated with tris(2-carboxyethyl)phosphine hydrochloride (TCEP) and iodoacetamide (IAA) to modify residual free cysteines. The resulting peptides were subjected to DIA-MS to identify and quantify cysteine-containing peptides. The percentage of oxidative modification was calculated by comparing the abundance of carbamidomethylated cysteine-containing peptides in each sample (Fig. 1A). Using this method, we investigated the effect of MND on the proteome redox status in DU145 cells. To identify the optimal concentration of MND for redox proteome analysis, we assessed the response of DU145 cells to various doses of MND by measuring intracellular ROS levels with CellROX™ and reduced cysteine levels with 5,5’-dithiobis(2-nitrobenzoic acid) (DTNB). At 50 µM MND, we observed a significant increase in intracellular ROS and a decrease in reduced cysteine levels (Fig. 1B, 1C). Based on these results, we selected 50 µM MND as the concentration for subsequent redox proteome analysis. The samples were prepared from four biological replicates, resulting in a dataset of 69,131 peptides (Sup. Table S1), which included 10,543 cysteine-containing peptides (Fig. 1D, Sup. Table S2). A total of 6812 peptides were quantified in all eight samples, in which all cysteines were modified with IAA. Among these, total 3671 peptides were quantified in all four DMSO- or MND-treated replicates, with 2459 peptides overlapping (Fig. 1D, Sup. Table S3).

**Figure 1.**
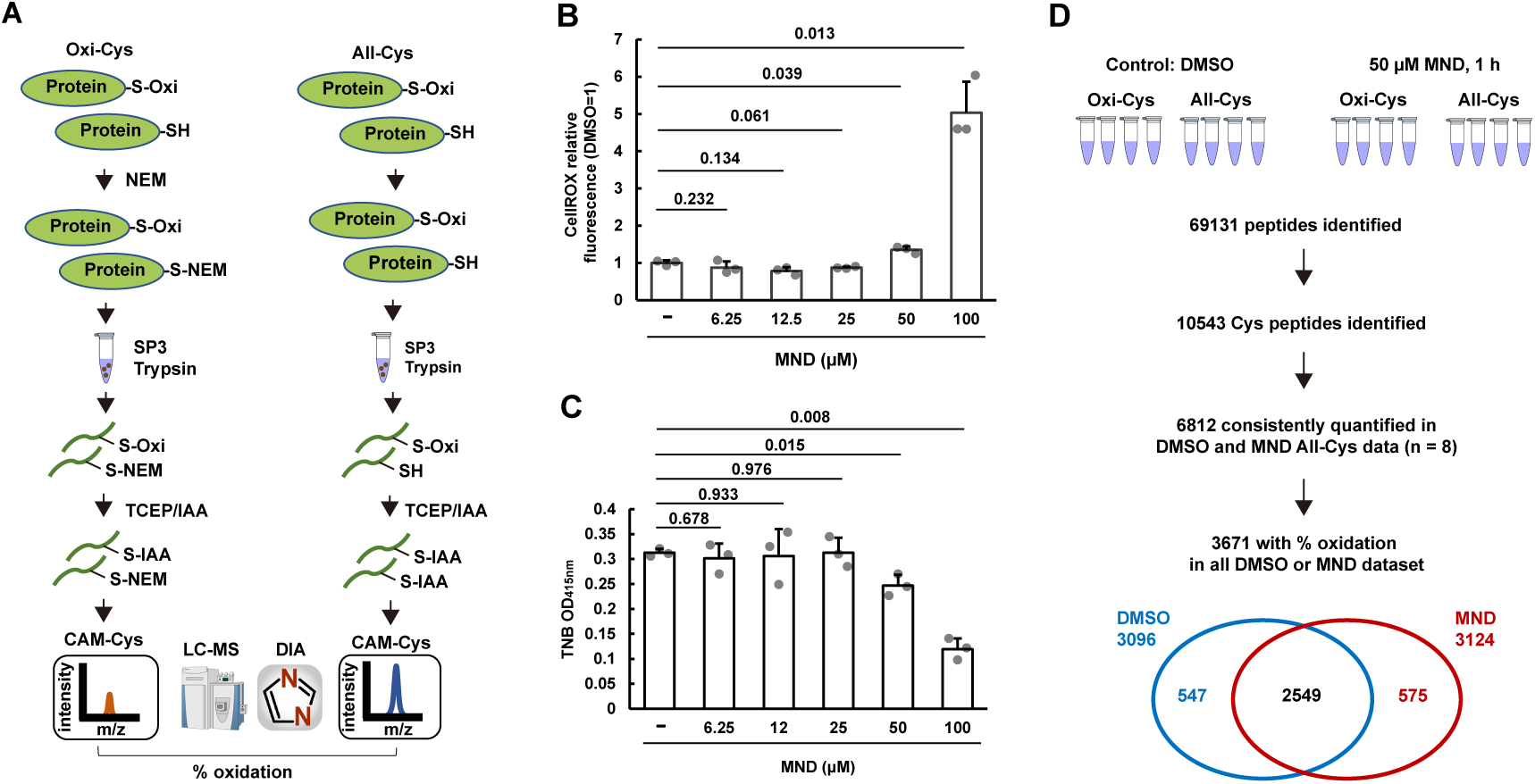
DIALRP application to the analysis of menadione response in DU145 cells. (A). Workflow of DIALRP. (B, C) Quantification of ROS using CellROX (B) and reduced cysteines using DTNB (C) in cells under the treatment of indicated concentrations of MND for 1 h. The data were shown in mean ± SD of 3 biological replicates represented as single points. Statistical significance was calculated using two-tailed paired t-test versus DMSO-treated (-), and the obtained *p*-values are indicated. (D) Schematic illustration of identified and quantified cysteine-containing peptides derived from the samples in DMSO- and 50 µM MND-treated cells.

Next, we validated the reliability of the DIALRP quantification data by comparing the oxidation percentages across four replicate samples from DMSO- and MND-treated cells (Sup. Table S4). The correlation coefficients for the oxidative modification ratios in the DMSO-treated samples ranged from 0.840 to 0.887 (Sup. Fig. S1). In the quantitative data for the DMSO and MND treatments, 74.0% and 72.1% of values, respectively, had a coefficient of variation (CV) of less than 0.2, with median CVs of 0.143 and 0.136 (Fig. 2A). The median percentage of oxidative modification in each data set averaged 24.3% for DMSO treatment (rep 1: 25.8%, rep 2: 22.8%, rep 3: 24.9%, rep 4: 23.8%) and 36.4% for MND treatment (rep 1: 37.2%; rep 2: 38.1%; rep 3: 34.7% %; rep 4: 35.6%) (Fig. 2B). A comparison of the distribution of the average percentage of oxidative modification for each peptide in the DMSO- and MND-treated samples showed a significant increase in the percentage of oxidative modification after MND treatment (Fig. 2C). Additionally, the MND treatment caused a significant increase in the percentage of oxidative modification in most peptides (Fig. 2C). These results confirm the robustness and reproducibility of DIALRP in quantifying cysteine oxidative modifications under different conditions.

**Figure 2.**
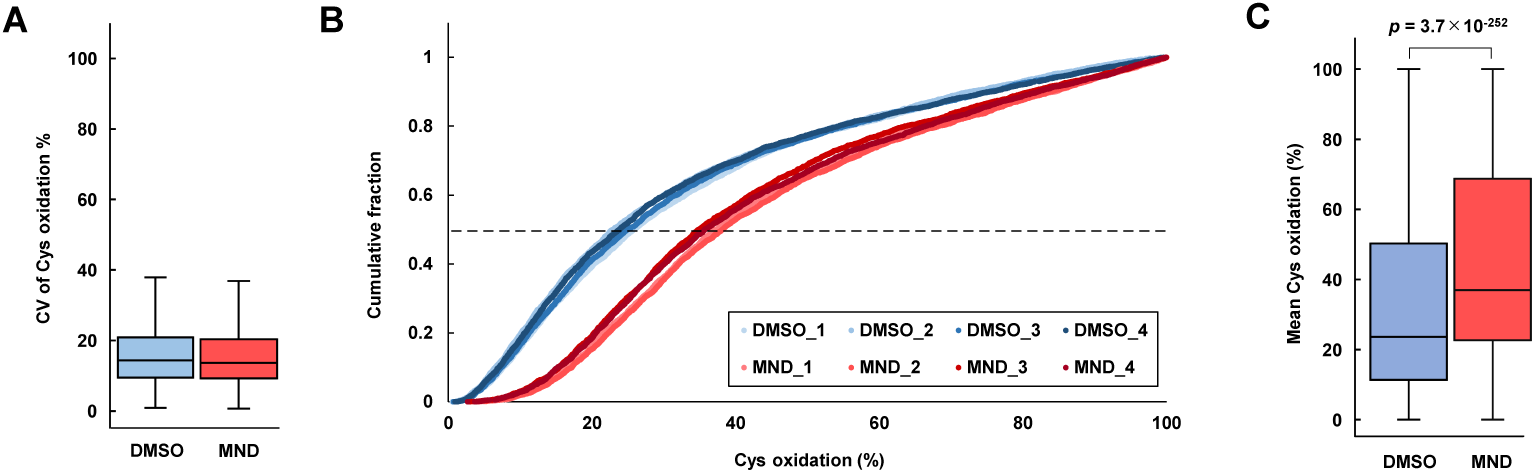
Validation of DIALRP quantitation results. (A) Boxplots of the coefficient of variation (CV) of cysteine oxidation percent in each peptide between four replicates of DMSO- and MND-treated DU145 cells. Boxplots display 25th and 75th percentile (bounds of box), median (center line), and largest and smallest value (whiskers) of the distribution. (B) Distribution of cysteine oxidation percent in each dataset of DMSO-treated and MND-treated cells. Dotted horizontal line (Y axis = 0.5) showed the median of each cysteine oxidation percent. (C) Boxplots of 3671 mean cysteine oxidation rate (%) in each peptide of DMSO-treated and MND-treated cells. Boxplots display 25th and 75th percentile (bounds of box), median (center line), and largest and smallest value (whiskers) of the distribution. Statistical significance was calculated using two-tailed paired t-test between DMSO-and MND-treated samples, and the obtained *p*-values are indicated.

As redox proteomics is primarily performed using stable isotopes with data-dependent acquisition mass spectrometry, enrichment or fractionation of peptides has been used to improve the number of identified peptides. This complicates the sample preparation for redox proteome analysis. To resolve this problem, label-free redox proteomics was reported in Redox Biology in 2019, which profiles the cysteine redox status based on DIA/SWATH(Anjo *et al*, 2019). As noted in this earlier report, the advantage of label-free analysis is that the use of non-isotope-labeled alkylation reagents, which are commonly used in most proteomics laboratories, incurs no cost during sample preparation. Additionally, label-free analysis has the advantage of fewer restrictions on the number of samples. In recent years, the use of DIA-MS for protein identification has increased dramatically with the development of software tools(Lou & Shui, 2024). Using the widely available Q-Exactive mass spectrometer, we successfully quantified redox changes in 3671 cysteine peptides with high reliability through a deep learning-based, library-free search using DIA-NN(Demichev *et al*, 2020). Several features of the approach contribute to its accuracy. First, it leverages DIA-MS, known for its precise and reliable quantitation. Second, unlike enrichment-based methods, which are prone to handling errors and complicated data correction, our method allows straightforward normalization using total peptide ion intensities, enhancing the reliability and consistency of quantitative results.

### DIALRP revealed that a group of thiols in translation-related proteins are most affected in response to menadione in DU145 cells

To evaluate the effect of MND on the oxidation state of cysteine, only cysteine-containing peptides were selected for comparison of quantitative values. Among the 1,467 cysteine-containing peptides with a q-value cutoff of less than 0.01, 1,017 showed a more than 2-fold change in oxidation state upon MND treatment (Fig. 3A, Sup. Table S4). The same set of eight samples, in which all cysteine residues were modified with IAA, was also analyzed to assess changes in protein expression independent of cysteine oxidation (Fig. 3B, Sup. Table S5). These data revealed that the cysteine redox response to MND treatment showed dramatic changes, whereas the overall protein expression levels remained relatively stable. Among the 1,467 MND-sensitive cysteine-containing peptides, the top 25% (367 peptides) were identified as highly responsive to MND treatment, corresponding to 295 proteins with strongly MND-sensitive cysteines (Fig. 3C).

**Figure 3.**
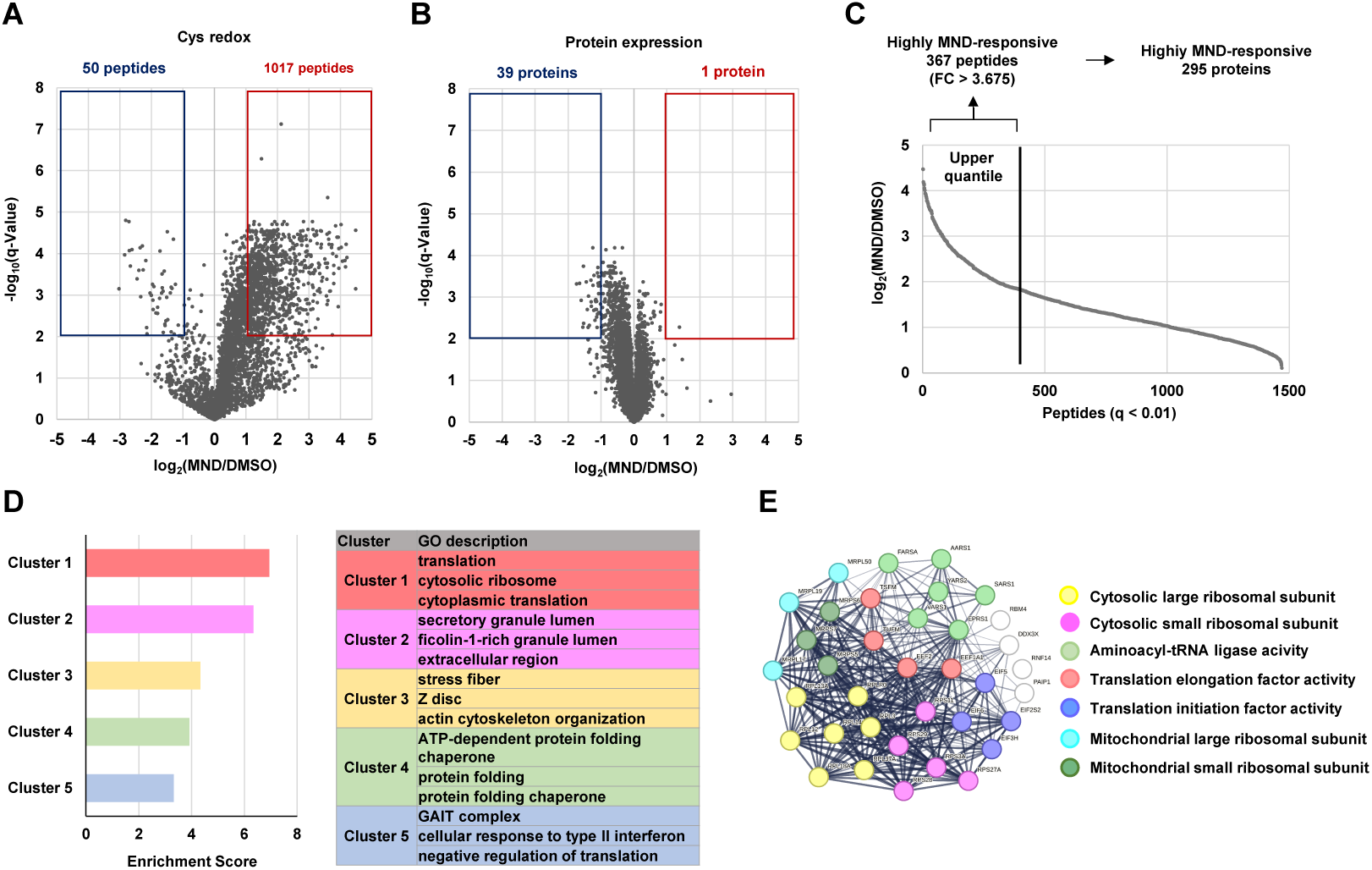
Extraction of significantly oxidized cysteines in response to MND and in silico prediction of their functions. (A, B) Volcano plot showing log_2_(fold change) in oxidation percentage for 3,671 unique cysteine-containing peptides (A) and expression of 6104 proteins (B) of MND-versus DMSO-treated cells plotted against −log_10_(*q-*value). (C) Order of 1,467 significantly oxidized cysteine peptides with a q-value cutoff of less than 0.01. The top 25% (367 peptides with fold change more than 3.68) were identified as highly responsive to MND treatment, corresponding to 295 proteins with strongly MND-sensitive cysteines. (D) GO analysis of 295 proteins with strongly MND-sensitive cysteines using DAVID functional annotation clustering. Enrichment scores for the top 5 clusters are shown in the bar graph (left panel), and the 3 GO descriptions with the lowest *p*-value in each cluster are shown in the table (right panel). (E) Functional molecular network of 36 highly MND-responsive cysteine-containing proteins annotated with GO “translation.” The 7 significantly enriched GO annotation selected in the list of functional enrichments in STRING were used to categorize these proteins.

Gene ontology (GO) enrichment analysis using DAVID revealed that the 295 proteins that were highly responsive to MND treatment were associated with various cellular processes, including translation (cluster 1), extracellular secretion (cluster 2), actin cytoskeleton organisation (cluster 3), chaperone activity (cluster 4), and interferon-responsive pathways (cluster 5) (Fig. 3D, Sup. Table S6). Among these categories, translation-related proteins were the most significantly enriched (cluster 1), indicating the strong impact of MND on translational mechanisms. Further analysis of the 36 proteins annotated with the GO term "translation" using STRING classified them into seven functional groups, such as cytosolic and mitochondrial ribosomal proteins, translation factors, and aminoacyl-tRNA synthetases (Fig. 3E, Sup. Table S7). These results suggest that various translation-related factors respond to oxidative stress and regulate their function at multiple stages of the translation process.

### Validation of cysteine redox response of translation-related proteins

Since the group of proteins most affected by MND-induced oxidative stress was thought to be translation-related, the translation status of MND-treated DU145 cells was evaluated by nascent polypeptide labelling using l-azidohomoalanine (AHA). We observed a decrease in translational activity with increasing concentrations of MND (Fig. 4A, 4B). Especially, translational activity was drastically decreased under conditions treated with 50 µM menadione, consistent with increased intracellular ROS and elevated cysteine oxidation (Fig. 1B, 1C). This decrease in translational activity was reversed by N-acetylcysteine (NAC) treatment (Fig. 4C). The redox statuses of seven translation-related molecules, RPL12, RPL11, EIF2S2, EIF6, EEF2, TSFM, and MRPL50, with particularly elevated oxidative modifications were selected for further validation of the redox proteome analysis data (Fig. 4D). The FLAG-tagged forms of these proteins were expressed in HeLa cells and analyzed by western blotting using the maleimide-PEG reagent. We used HeLa cells for these validations because of the high transfection efficiency to sufficiently express all constructs, and confirmed that the cysteines of endogenous EIF6 was oxidized in response to MND treatment in HeLa cells, as well as DU145 cells (Sup. Fig. S2). The results confirmed that these proteins underwent oxidative modification in response to menadione, and that NAC treatment restored the redox status (Fig. 4E). These data support the redox proteomic data of the MND treatment response.

**Figure 4.**
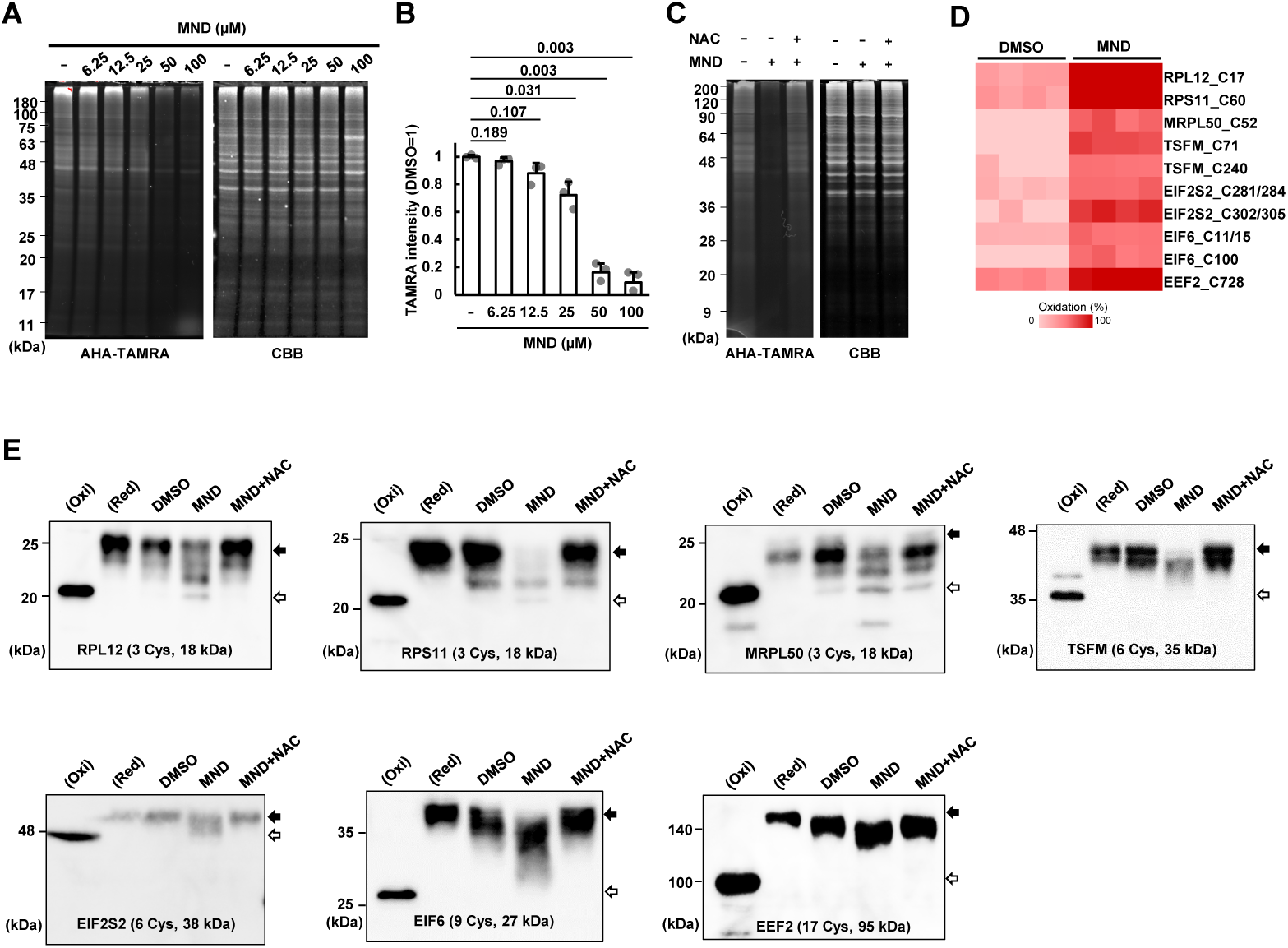
Validation of MND-induced oxidative stress response of a group of translation-related factors. (A, B) Quantification of nascent polypeptides in DU145 cells under the treatment of indicated concentrations of MND for 1 h using l-azidohomoalanine (AHA). (A) Total proteins were separated by SDS-PAGE, and AHA-tetramethylrhodamine (TAMRA)-labeled nascent polypeptides were detected by fluorescent scanner. Total proteins were visualized by CBB staining. Representative images of AHA-TAMRA and CBB were shown. The fluorescent intensities were quantified and normalized as 1 for DMSO-treated samples (B). The data were shown in mean ± SD of 3 biological replicates represented as single points. Statistical significance was calculated using two-tailed paired t-test versus DMSO-treated (-), and the obtained *p*-values are indicated. (C) Detection of nascent polypeptides in DU145 cells treated with DMSO, 50 µM MND or 50 µM MND plus 5 mM N-acetylcysteine (NAC) for 1 h using AHA. (D) Redox proteomics data of peptides with particularly elevated oxidative modifications in selected 7 translation-related proteins. (E) Validation of MND-oxidative stress response by redox western blotting. HeLa cells transfected with plasmid expressing each FLAG-tagged protein were treated with DMSO, 50 µM MND or 50 µM MND plus 5 mM NAC. The cell lysate was treated with mPEG-maleimide and analyzed. The proteins treated with TCEP and NEM (mimicking fully oxidized form, Oxi), and TCEP and mPEG-maleimide (mimicking fully reduced form, Red) were detected simultaneously. Black and white arrows show fully oxidized and reduced forms, respectively.

### Functional validation of the translation factors under MND-induced oxidative stress conditions

To assess the functional influence of translation factors under oxidative conditions induced by MND treatment, changes in their subcellular localization and protein binding were analyzed. Of the seven translation-related factors evaluated for oxidative stress response by western blotting, the functions of three proteins, EIF2S2, EIF6, and EEF2, were successfully evaluated. FLAG-IP-MS analysis of EIF2S2, EIF6, and EEF2 predominantly identified proteins that were previously reported to interact physically and functionally, such as EIF2S1, EIF2S3, and CDC123 in EIF2S2; BCCIP and RPL23 in EIF6; and HGH1 in EEF2 (Fig. 5A, 5E, 5I, Sup. Table S8-S11).

**Figure 5.**
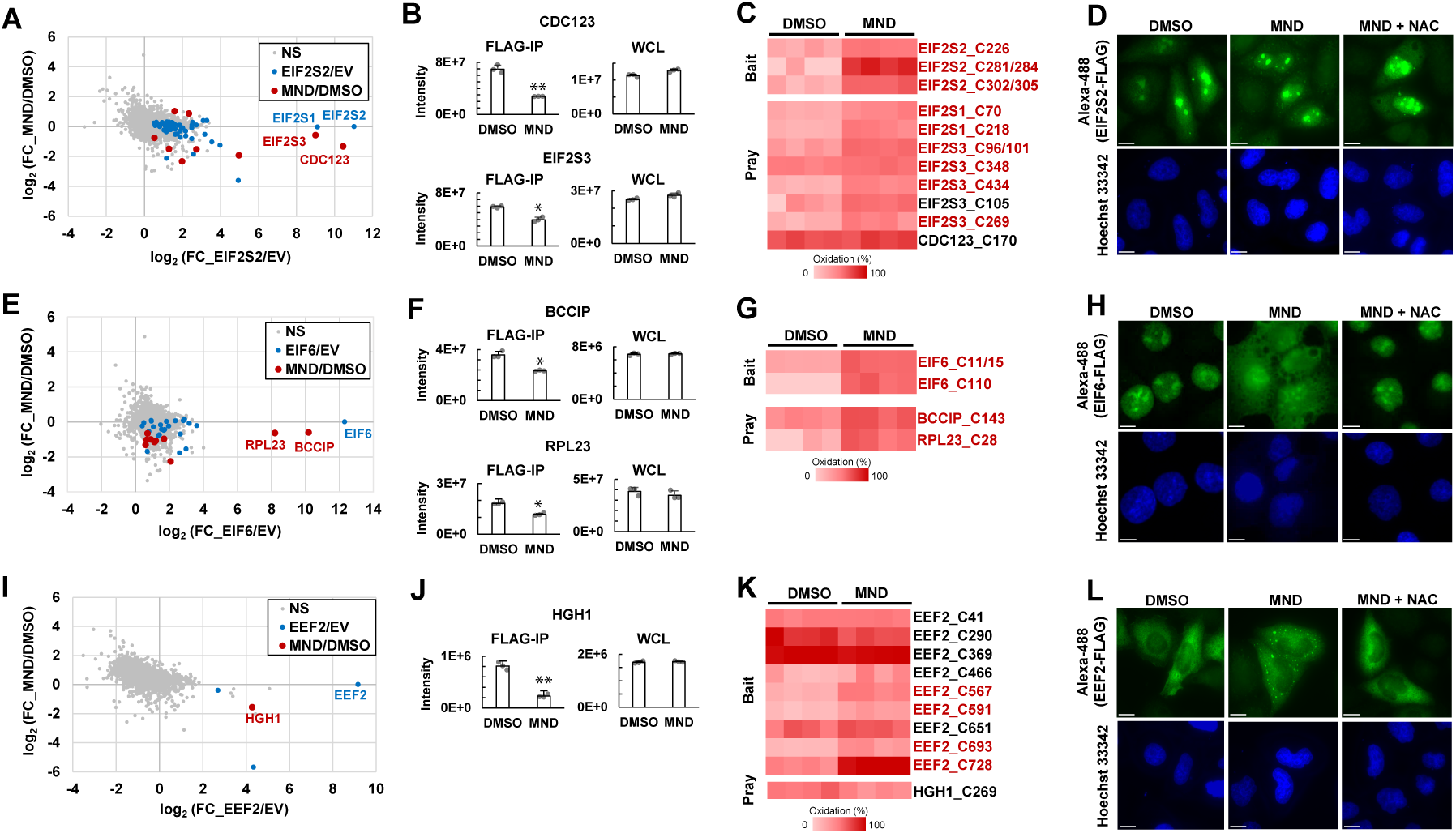
Functional validation of translation factors under MND treatment by FLAG-IP-MS and immunocytochemistry. (A, E, I) Scatterplots showing quantitative data of FLAG-IP MS analysis of EIF2S2 (A), EIF6 (E), and EEF2 (I). X-axis and Y-axis represent log_2_ fold change of intensities of proteins in FLAG-IP fraction prepared from HeLa cells transfected with bait-expressing plasmid compared to that with empty vector (EV) and in proteins in FLAG-IP fraction prepared from bait-expressing cells treated with 50 µM MND for 1 h compared to those treated with DMSO, respectively. Proteins that were significantly more abundant in the bait FLAG-IP fraction compared to EV are indicated by blue dots. Of those proteins, their bindings significantly altered by MND treatment are indicated by red dots. The names of baits, and proteins that have been reported to form complexes, are listed in the graph. (B, F, J) Bar graphs showing the intensities of EIF2S2 (B), EIF6 (F), and EEF2 (J) binding proteins to significantly altered by MND treatment in FLAG-IP fraction and whole cell lysate (WCL). The data were shown in mean ± SD of 3 biological replicates represented as single points. \**q* < 0.05, \*\**q* < 0.01 versus DMSO-treated (-). Each *q*-value is listed in Sup. Table S9-S11. (C, G, F) Redox proteomics data of the EIF2S2 (C), EIF6 (G), and EEF2 (K) and their binding proteins. Cysteines that were oxidized significantly with a q-value less than 0.01 was highlighted in red. (D, H, L) Fluorescent images showing the localization of EIF2S2 (B), EIF6 (F), and EEF2 (J) in HeLa cells treated with DMSO, 50 µM MND or 50 µM MND plus 5 mM NAC. The cells were counterstained with Hoechst33342 to detect their nuclei. Scale bar = 10 µm.

We observed a decrease in the binding of EIF2S3 and CDC123 to EIF2S2 in response to MND treatment (Fig. 5A, 5B), suggesting that formation of the eIF2 complex is inhibited by MND-induced oxidative stress. Immunohistochemistry revealed no change in EIF2S2 localization, which remained primarily in the nucleoli despite MND treatment (Fig. 5D). We also observed a decrease in the binding of BCCIP and RPL23 to eIF6 (Fig. 5E and 5F). Previous studies reported that eIF6, BCCIP, and RPL23 form a tripartite complex, with BCCIP functioning to translocate eIF6 to the nucleus and nucleoli(Wyler *et al*, 2014; Ye *et al*, 2020). Although eIF6 was primarily localized in the nucleus and nucleolus, even in its FLAG-tagged form, MND treatment markedly increased its cytoplasmic localization. This shift was reversed by simultaneous treatment with MND and NAC, which restored eIF6 localization in the nucleus and nucleolus (Fig 5H). Moreover, MND treatment decreased the binding of HGH1 to EEF2 (Fig. 5I, 5J), an interaction in which HGH1 functions as a chaperone of EEF2(Mönkemeyer *et al*, 2019; Schopf *et al*, 2019). Immunocytochemistry revealed that EEF2-containing aggregates formed in the cytoplasm in response to MND treatment, but dissipated with concurrent NAC treatment (Fig. 5L). Notably, quantitative DIA-MS analysis of cell lysates confirmed that the observed changes in protein interactions and localization after MND treatment occurred independent of any changes in protein expression levels (Fig. 5B, 5F, 5J, Sup. Table S15).

These results indicated that specific cysteine residues within translation-related factors are particularly susceptible to oxidative modifications under MND-induced stress, with potential functional implications. In the EIF2 complex, 8 of the 11 cysteine-containing peptides (covering 10 of 14 cysteines) in EIF2S1, EIF2S2, EIF2S3, and CDC123 were significantly oxidatively modified (Fig. 5C). This suggests that oxidative modifications affect the function of the eIF2 complex, possibly affecting the initiation of translation. In the eIF6-BCCIP-RPL23 ternary complex, all four detected peptides (with five cysteines) were significantly oxidatively modified (Fig. 5G), indicating that oxidative stress disrupted the normal localization and function of eIF6 in translational regulation. In contrast, EEF2 showed oxidative modification in four of its nine cysteine-containing peptides, whereas HGH1, a known chaperone of EEF2, displayed no detectable oxidative modifications (Fig. 5K). This difference suggests a protective role for HGH1 under oxidative conditions, potentially helping stabilize EEF2.

The FLAG-IP-MS analysis of two proteins, RPL12 and MPRL50, failed to significantly identify binding proteins, despite the high abundance of identified baits; the analysis of TSFM was able to identify TUFM as a binding protein, but MND treatment had no significant effect on binding (Sup. Fig. S3, Sup. Table S12-S14). In the case of RPS11, enrichment by FLAG-IP was not successful, and did not reach a sufficient level for analysis. Immunocytochemistry was performed to evaluate whether their localization was altered in response to the MND treatment (Sup. Fig. 4). FLAG-tagged RPL12 is mainly localized to the nucleolus. RPL12 localization to the nucleolus was disrupted by MND treatment, suggesting that normal RPL12-related ribosomal biogenesis may be inhibited by oxidative stress. We also observed that the localization of RPS11, TSFM, and MRPL50 was not altered by MND treatment.

Overall, these findings highlight that MND-induced oxidative stress selectively affects cysteine residues in translation-related factors, potentially modulating translation as part of the adaptive response of cells to oxidative stress.

Oxidative modification of cysteine has been reported to regulate various biological processes such as signal transduction(Heppner *et al*, 2018), metabolic adaptation(Van Der Reest *et al*, 2018), and developmental hematopoiesis(Pimkova *et al*, 2022). Hence, methods for quantifying the oxidative modification of cysteine are widely applicable. Current redox proteome analysis techniques require relatively complex sample-handling methods. We have demonstrated a workflow that enables redox proteome analysis using an approach comparable to sample handling in general expression proteome analysis. Translation-related factors were the most significant group of proteins that underwent oxidative modifications during MND-induced oxidative stress. Previous redox proteomic studies have also reported enhanced oxidative modification of translation-related factors in response to oxidative stress, indicating that translation and cysteine redox are closely related(Go *et al*, 2013; Topf *et al*, 2018). Indeed, the decrease in the translational activity of DU145 cells during MND treatment was accompanied by a significant increase in ROS and a decrease in intracellular reduced cysteine levels (Fig. 1A, 1B, 4A, 4B). The accompanying oxidative response was particularly pronounced in a group of factors that play a central role in translation initiation and elongation, such as EIF2, eIF6, and EEF2 (Fig. 5). Previous studies reported that the functions of eIF2, eIF6, and eEF2 are suppressed under the stress condition through phosphorylation by kinases: eIF2α by PERK, GCN2, PKR, and HFI(Wek *et al*, 2006), eIF6 by RACK1 and GSK3β(Ceci *et al*, 2003; Jungers *et al*, 2020), and eEF2 by eEF2K(Ryazanov *et al*, 1988; Liu & Proud, 2016). A previous report showed that menadione treatment causes increased phosphorylation of Ser51 of eIF2α and decreased translational activity, whereas inhibition of eIF2α phosphorylation by PERK inhibitors does not restore the decreased translational activity(Samluk *et al*, 2019). These findings suggest the existence of a sophisticated regulatory network in which crosstalk between cysteine oxidation, phosphorylation, and other post-translational modifications orchestrates translational control during oxidative stress. Further studies are required to elucidate the intricate details of these multifaceted regulatory mechanisms.

The SP3 method (Hughes *et al*, 2019) significantly enhanced the throughput and scalability of the DIALRP workflow, making high-throughput redox proteomic analysis more feasible. Unlike traditional methods that often require complex and time-consuming steps for cysteine peptide enrichment or stable isotope labeling, the SP3 approach streamlines sample preparation and enables efficient parallel processing of multiple samples(Hughes *et al*, 2019). This streamlined approach is well suited for large-scale studies, such as those investigating oxidative stress across various cell types or conditions. The high-throughput capability of DIALRP with the SP3 method accelerates redox proteomics research, enabling comprehensive cysteine oxidation profiling across biological and clinical contexts, and providing a valuable tool for broader redox biology studies.

In summary, this study provides a comprehensive framework for the analysis of cysteine oxidative modifications using DIA-MS-based label-free redox proteomics, specifically revealing the oxidative stress response in DU145 prostate cancer cells. By identifying translation-related factors as primary targets of oxidative modification, we highlighted a potential mechanism by which oxidative stress modulates protein synthesis and cellular adaptation. However, this study was limited to the oxidative stress response to menadione treatment in DU145 cells, which are highly resistant to oxidative stress(Jayakumar *et al*, 2014; Tossetta *et al*, 2023). Although we confirmed the oxidative stress response of selected translation-related factors when they were overexpressed in HeLa cells, this was not confirmed in other cell types or under other oxidative stress conditions. Therefore, a detailed analysis of the differences in oxidative stress tolerance and oxidative stress conditions induced by DIALRP will lead to a better understanding of the oxidative stress response in cancer cells.

## Materials and Methods

### Cell line and culture conditions

DU145 cells were cultured under 5% CO_2_ at 37 °C in RPMI1640 medium (Fujifilm-Wako, Japan) supplemented with 10% fetal bovine serum (FBS) (NICHIREI BIOSCIENCES, Japan), Penicillin/streptomycin (10,000 U/mL) (Thermo Fisher Scientific, MA, USA), and Sodium pyruvate (100 mM) (Thermo Fisher Scientific, MA, USA). HeLa cells were cultured under 5% CO2 at 37 °C in Dulbecco’s modified Eagle’s medium (Fujifilm-Wako, Japan) supplemented with 10% FBS (NICHIREI BIOSCIENCES, Japan), Penicillin/streptomycin (10,000 U/mL) (Thermo Fisher Scientific, MA, USA), MEM Non-Essential Amino Acids Solution (100×) (Thermo Fisher Scientific, MA, USA), and Sodium pyruvate (100 mM) (Thermo Fisher Scientific, MA, USA).

### Cell lysate preparation

For reduced cysteines measurement by DTNB, redox proteomics, and redox western blotting, cells were washed with PBS twice and solubilized in lysis buffer containing 7 M Urea, 1% Triton X-100, 100 mM HEPES-NaOH (pH 7.2) for 15 min at 4 °C on a shaker. For FLAG-IP-MS analysis, cells were washed with PBS twice and solubilized in lysis buffer containing 50 mM Tris-HCl (pH 7.4), 150 mM NaCl, 0.5% Triton X-100, 5 mM MgCl_2_, and 1% (v/v) protease inhibitor mixture (Roche, Switzerland) for 15 min at 4 °C on a shaker. Lysates were centrifuged at 20,000 × g for 15 min at 4 °C, and the protein concentration of the supernatants was determined using the BCA Protein Assay Kit (Takara Bio, Inc., Japan).

### ROS measurement

After treated with menadione, culture media was removed, and the cells were incubated with the culture media containing 5 μM CellROX^TM^ Green Reagent (Thermo Fisher Scientific, MA, USA) and 0.5 μg /ml Hoechst33342 (Dojindo, Japan) at 37 °C for 30 min. Cells were collected using trypsin, washed with PBS, and gently resuspended in PBS. The fluorescence intensities of CellROX and Hoechst were measured using a Fluoroskan FL microplate reader (Thermo Fisher Scientific).

### Measurement of reduced cysteines in cell lysate

Fifty μg of proteins were treated with 200 μM DTNB (Dojindo, Japan) in 50 μl of lysis buffer for 10 min at room temperature. The absorbances of in 5-Mercapto-2-nitrobenzoic acid (TNB) in the samples was measured at 415 nm using a BioRad iMark microplate reader (Bio-Rad Laboratories, TX, USA).

### Sample preparation for redox proteome analysis

Ten μg of proteins were treated with 10 mM NEM in 50 μl of lysis buffer. The NEM was eliminated using the SP3 method(Hughes *et al*, 2019). For the sample to quantify all of cysteine peptides, lysates containing 10 μg of proteins were directly applied to SP3 beads. SP3 beads were suspended with 50 μl of 100 mM HEPES-NaOH (pH 8.0) containing 0.4 μg of trypsin (Richcore, India), and trypsin digestion was performed at 37 °C for 16 h. The supernatant was collected and treated with 2.5 mM TCEP for 30 min at 37 °C, followed by treatment with 10 mM IAA for 30 min at room temperature. Trifluoroacetic acid (TFA) was added to the samples at final concentration of 1%. The sample was desalted using styrenedivinylbenzene (SDB)-StageTip (Rappsilber *et al*, 2007) and dissolved in 0.1% TFA in 2% acetonitrile for LC-MS analysis.

### Plasmid construction and transfection

To obtain cDNA, total RNA was extracted from HeLa cells using TRIzol reagent (Thermo Fisher Scientific) according to the manufacturer’s protocol. Reverse transcription of first strand cDNA was performed with ReverTra Ace^®^ (TOYOBO, Japan) using the total RNA as a template. The PCR was performed with KOD One^TM^ PCR Master Mix (TOYOBO, Japan) The forward and reverse primers for RPL12, RPS11, MRPL50, TSFM, EIF2S2, EIF6, EEF2 were designed to ligate these cDNAs with expression plasmid as follows: CAGGCCCGAATTCGGTCCACCATGCCGCCGAAGTTCGACC (RPL12 forward), TCCTTGTAGTCGCCGGTACCTCGAGAACTGGCTGGGCATTCCACAGCACCAC (RPL12 reverse), CAGGCCCGAATTCGGGGGAAGATGGCGGACATTCAGACTG (RPS11 forward), TCCTTGTAGTCGCCGGTACCTCGAGAGAACTTCTGGAACTGCTTCTTGGTGC (RPS11 reverse), CAGGCCCGAATTCGGTCGAAGATGGCGGCGCGATCTGTGTC (MRPL50 forward), TCCTTGTAGTCGCCGGTACCTCGAGAGTAACTCCAAGTGATTTTCAAATTGGG (MRPL50 reverse), CAGGCCCGAATTCGGAGAGAGATGTCGCTGCTGCGGTCG (TSFM forward), TCCTTGTAGTCGCCGGTACCTCGAGATTCAGTTTCTGCTGCCTCTTCACCTTC (TSFM reverse), CAGGCCCGAATTCGGGCAGCCATGTCTGGGGACGAGATG (EIF2S2 forward), TCCTTGTAGTCGCCGGTACCTCGAGAGTTAGCTTTGGCACGGAGCTGTGCTC (EIF2S2 reverse), CAGGCCCGAATTCGGATGGCGGTCCGAGCTTCGTTCGAG (EIF6 forward), TCCTTGTAGTCGCCGGTACCTCGAGAGGTGAGGCTGTCAATGAGGGAATC (EIF6 reverse), CAGGCCCGAATTCGGGCCACCATGGTGAACTTCACGGTAG (EEF2 forward), TCCTTGTAGTCGCCGGTACCTCGAGACAATTTGTCCAGGAAGTTGTCCAGGG (EEF2 reverse), respectively. The PCR was performed using a Bio-Rad T100 Thermal Cycler (Bio-Rad Laboratories, TX, USA). with the following thermal cycling parameters: 94 °C for 2 min, followed by 35 cycles of 98 °C for 15 s, 56 °C for 30 s, and 72 °C for 1 min, and the final extension was performed at 72 °C for 10 min. The PCR products were cloned into pCMV-(DYKDDDDK)-C vector (Clontech, CA, USA) using Gibson Assembly^®^ Master Mix (NEB, MA, USA) according to manufacturer’s protocol, and were sequenced. The constructed expression plasmids were used to transfect the cultured cells. HeLa cells were transfected with expression plasmids using Fugene HD reagent (Promega, Madison, WI, UAS) according to the manufacturer’s protocol.

### FLAG-Immunoprecipitation and sample preparation for MS analysis

Five μl of anti-FLAG M2 Magnetic Beads (Sigma-Aldrich, MO, USA) were washed twice with 200 μl of TBS (50 mM Tris-HCl, 150 mM NaCl, pH 7.4) buffer. The cell lysate containing 100 μg of protein amount was added to the washed resin beads. The mixture was incubated for 30 min at 4 °C with gentle shaking. The beads were washed three times with 400 μl of TBS. To prepare the sample for mass spectrometry, 50 μl of 0.5 mg/ml FLAG solution in 50 mM Tris HCl pH 7.4, 150 mM NaCl, 0.05 % Triton X-100 were added to the washed resin beads, and the mixture was incubated at room temperature with gentle shaking for 5 min. The supernatant was obtained and further treated to analyze. One hundred sixty microliters of solution containing 8.75 M urea and 50 mM Tris-HCl (pH 8.5) were added to 40 µL of eluted sample. Samples were treated with 2.5 mM TCEP for 30 min at 37 °C, followed by treatment with 10 mM IAA for 30 min at room temperature. Acetone was added to the sample at final concentration of 80%, mixed, and incubated for 30 min at room temperature. The sample was centrifuged at 20, 000 ×*g* for 15 min at room temperature to precipitate proteins. The precipitate was washed twice with 1 ml of 90% acetone. After dried, 50 µL of 100 mM HEPES-NaOH pH 8.0 containing 200 ng trypsin were added and incubated at 37 ℃ for 16 h. TFA was added to the sample at final concentration of 1%. The sample was desalted using SDB-StageTip (Rappsilber *et al*, 2007) and dissolved in 0.1% TFA and 2% acetonitrile for LC-MS analysis. Triplicate biological samples were prepared from all the bait samples. Triplicate biological samples from HeLa cells transfected with the empty vector were prepared as controls to extract the binding proteins. To evaluate the change of expression of the binding proteins, HeLa cells treated with 50 µM MND for 1 h were lysed with the same solution using FLAG-IP-MS analysis. Forty µL of solution containing 8.75 M urea and 50 mM Tris-HCl (pH 8.5) were added to 10 µl of cell lysate containing 10 µg of proteins. Samples were treated with 2.5 mM TCEP for 30 min at 37 °C, followed by treatment with 10 mM IAA for 30 min at room temperature. The sample was applied to SP3 methods (Hughes *et al*, 2019) to remove TCEP and IAA, suspended with 50 μl of 100 mM HEPES-NaOH (pH 8.0) containing 0.4 μg of trypsin (Richcore, India), incubated at 37 °C for 16 h. TFA was added to the sample at final concentration of 1%. The sample was desalted using SDB-StageTip (Rappsilber *et al*, 2007) and dissolved in 0.1% TFA and 2% acetonitrile for LC-MS analysis.

### LC-MS

Peptide samples were injected into a pre-column (L-column2 micro, CERI, Japan) and fractionated on an in-house fabricated 15-cm column packed with 2-μm octadecyl silane particles (CERI, Japan). Elution was performed with a linear gradient of 5%−32% solvent B over 60 min at a flow rate of 200 nl/min (solvent A = 0.1% formic acid; solvent B = 0.1% formic acid in acetonitrile) with the use of a Dionex Ultimate 3000 HPLC System (Thermo Fisher Scientific, MA, USA). The eluted peptides were sprayed with a nanoelectrospray source and a column oven set at 42 °C (AMR, Japan). The Q Exactive Hybrid Quadrupole-Orbitrap mass spectrometer (Thermo Fisher Scientific, Waltham, MA, USA) was operated in DIA mode. All data were acquired in the profile mode using positive polarity. MS1 spectra were collected in the *m/z* range of 430–860 at a resolution of 35,000 using an AGC target value of 1 × 10^6^ with a maximum injection time of 50 ms. MS2 spectra were collected in the range of > 200 *m/z* at a resolution of 17,500 using an AGC target value of 1 × 10^6^ and automatic maximum ion ITs. Twenty-one DIA windows of 20 units ranged from 430 to 850 *m/z*, with an overlap of one unit. The normalized collision energy was set to 25.

### MS data processing

All raw DIA data were processed using DIA-NN (ver. 1.8.1) (Demichev *et al*, 2020) using a library-free search mode against human UniProt/SwissProt sequences (release 2024_02, 20434 entries) to identify tryptic peptides. The enzyme specificity was set to ‘trypsin’. Up to one missed trypsin cleavages were allowed. Carbamidomethylation of cysteine was set as a fixed modification. FDR threshold for peptide identification was set to 1%. Match between run mode was used for both DIALRP and FLAG-IP MS. For DIALRP and FLAG-IP MS, cross-run normalization was set as ‘RT-dependent and signal dept.’ and ‘None’, respectively. For DIALRP, the positions of the cysteines in all identified peptides were annotated based on UniProt/Swiss-Prot sequences using a Python script. The oxidation percentage was calculated based on the intensity of carbamidomethylated cysteine-containing signals from NEM-treated (Oxi-Cys) and non-NEM-treated (All-Cys) samples. The oxidation percentages were averaged when redundantly quantified due to the difference in charges. The data with more than 100 % oxidation were defined as 100 %.

The significantly oxidized cysteine peptides, differentially expressed proteins, and altered binding proteins were extracted by statistical tests using Perseus software(Tyanova *et al*, 2016). For the analysis of oxidized cysteine peptides, missing oxidation percentages of peptides quantified in all four DMSO- or MND-treated replicates were applied to imputations from a normal distribution. Multiple testing with Student’s t-test and Benjamini-Hochberg FDR, with *q*-value of less than or equal to 0.01 as cut-off value, was performed to extract peptides that underwent significant oxidative modifications and differentially expressed proteins. For the analysis of FLAG-IP-MS data, the quantitation data of all replicates in the bait samples were normalized according to the bait intensities. The missing quantification values of proteins in all three bait samples treated with DMSO-treated were applied to imputations from a normal distribution. To extract binding proteins under DMSO-treated conditions and altered binding proteins under MND-treated conditions, multiple testing with Student’s t-test and Permutation-based FDR was performed with *q*-value less than or equal to 0.05 as the cut-off value.

### In-silico gene ontology (GO) enrichment and network analysis

GO enrichment analysis was performed by functional annotation clustering using DAVID(Huang *et al*, 2009; Sherman *et al*, 2022). The background was set to the whole Homo sapiens genome. GOTERM_BP_DIRECT, GOTERM_CC_DIRECT, and GOTERM_MF_DIRECT were selected for the analysis. All proteins, which were annotated with GO ‘translation (Accession: GO:0006412)’ were extracted and the network of these proteins was visualized using STRING (Szklarczyk *et al*, 2023) Ver 12.0 (https://string-db.org/). The parameters were set as follows: network type: full STRING network; meaning of network edges: evidence; active interaction sources: text mining, experiments, databases, co-expression, neighbourhood, gene function, and co-occurrence; minimum required interaction score: 0.400; maximum number of interactors to show: first shell = no more than 10 interactions; second shell = none. Functional classification was performed using enriched GO terms in the list of functional enrichment in STRING.

### Nascent polypeptide detection

The cells were washed once with PBS and incubated in methionine-free medium containing 100 μM L-azidohomoalanine (AHA) reagent (Biosynth, UK) at 37 °C for 2 h. The cells lysed with 50 mM Tris-HCl (pH 7.4), 150 mm NaCl, 0.5% Triton X-100, and 1% (v/v) protease inhibitor mixture (Roche, Switzerland), for 15 min at 4 °C on a shaker. Lysates were centrifuged at 20,000 × g for 15 min at 4 °C, and the protein concentration of the supernatants was determined using the BCA Protein Assay Kit (Takara Bio, Inc., Japan). Ten μg of proteins were treated with 10 μM tetramethylrhodamine (TAMRA)-alkyne (Sigma-Aldrich, MO, USA), 1 mM TCEP, 100 μM tris[(1-benzyl-1H-1,2,3-triazol-4-yl)methyl] amine (TBTA) and 1 mM CuSO_4_ in lysis buffer for 2 h at room temperature. The samples were precipitated with acetone to final concentration of 80% (v/v). Samples were centrifuged to precipitate the proteins at 20,400 × g for 15 min at 4 °C. The obtained precipitants were washed twice with 90% (v/v) acetone, and dissolved in SDS-PAGE sample buffer with the incubation at 95 °C for 5 min. All proteins were separated on 12 % SDS-polyacrylamide gels. The TAMRA-labeled nascent polypeptides were detected using a Bio-Rad ChemiDoc Touch MP imaging system (Bio-Rad Laboratories, TX, USA). TAMRA fluorescence intensity was quantified using ImageJ software(Schneider *et al*, 2012). To normalize the protein amounts, the gels were stained with Coomassie Brilliant Blue G-250 after TAMRA detection and reimaged using the same system.

### Redox western blotting

The lysate was treated with 10 mM mPEG-maleimide 1 K (Biopharma PEG Scientific Inc., MA, USA) in lysis buffer for 1 h at room temperature. For validation of EEF2, mPEG-maleimide 2 K (Biopharma PEG Scientific Inc., MA, USA) was used to analyze. To mimic fully oxidized or reduced forms, the lysate was treated with 2.5 mM TCEP for 30 min at 37 °C, followed by treatment with 10 mM NEM or mPEG-maleimide. After the treatment, the sample was precipitated with acetone at final concentration of 80% (v/v). Sample was centrifuged to precipitate the proteins at 20,000 × g for 15 min at 4 °C. The precipitant was washed twice with 90% (v/v) acetone, and dissolved in SDS-PAGE sample buffer with the incubation at 95 °C for 5 min. Ten µg of labelled proteins were separated onto 8, 12 or 14% SDS-polyacrylamide gels according to their molecular weights, transferred onto a PVDF membrane by electroblotting, and subjected to immunoblotting with the primary anti-mouse anti-DDDDK-tag antibody (MBL Life science, M185-3L, Japan) or anti-rabbit anti-eIF6 antibody (Cell Signaling Technology, 3833, MA, USA). After incubation with the primary antibody, the membrane was probed with a horseradish peroxidase-conjugated mouse secondary antibody (Promega). The image was visualized with SuperSignal™ West Pico PLUS Chemiluminescent Substrate (Thermo fisher Scientific, MA, USA) and was detected using a BioRad ChemiDoc Touch MP imaging system (Bio-Rad Laboratories, TX, USA).

### Immunocytochemical analysis

HeLa cells cultured on glass plates were fixed with 3.7% formaldehyde in PBS for 15 min at room temperature and then permeabilized with 0.1% Triton X-100 in PBS on ice for 15 min. After washing with PBS, cells were incubated in primary Anti-DDDDK antibody (Cell Signaling Technology, 14793, MA, USA) diluted in PBS containing 5% bovine serum albumin followed by anti-rabbit Alexa Fluor® 488-conjugated IgG (Invitrogen, CA, USA) for 60 min at room temperature. After washing with PBS, the glass plates were mounted on slide glasses using ProLong™ Glass Antifade Mountant with NucBlue™ Stain (Invitrogen, CA, USA). Images were acquired using a DMi8 microscope equipped with the Leica Thunder Imaging System (Leica, Germany). Images were processed using Leica LAS X software (Leica, Germany).

### Statistical analysis

All values are expressed as the mean with the ± S.D., and significant differences between groups were assessed using Student’s *t* test, except for MS data.

## Acknowledgments

We thank the staff of the Department of Omics and Systems Biology at Niigata University, especially Kiyotaka Oshikawa, Atsushi Hatano, Yoshitomi Kanemitsu, Ayaka Kakihara, Naomi Hatanaka, Takako Ichihashi, and Yoko Sawaguchi, for their helpful support. We are also grateful to the staff members of the Center for Research Promotion, School of Medicine, Niigata University Medical School for their important contributions to the experiments. This work was supported by MEXT/JSPS KAKENHI under Grant No. JP 20H05930 (M.M.), 22H02607(M.M.), JP22K07144 (D.K.); AMED-CREST under Grant No. 21gm1410006h0001 (M.M.); and GteX Program Japan Grant Number JPMJGX23B3 (M.M.).

## Author contributions

D. K. and M. M. designed and performed the experiments. D.K. collected data. D.K. and M.M. prepared the manuscript. T. T. contributed to the bioinformatics tools.

## Data availability

All raw MS data were stored in jPOSTrepo (https://repository.jpostdb.org/)(Okuda *et al*, 2017). jPOSTIDs/PXIDs for the projects containing these data were JSPT003469/PXD057921 (DIALRP), JSPT003470/PXD057922 (FLAG-IP-MS), and JSPT003471/PXD0579230 (Expression proteome analysis of MND response in HeLa cells).

## Figure Legends

**Supplementary Figure S1 (related to Figure 2)**

Correlation coefficients of cysteine oxidation rates between each dataset and the four replicates of DMSO-treated samples. Scatter plots of the cysteine oxidation percentages of each peptide and their correlation coefficients are shown in the upper and lower boxes, respectively.

**Supplementary Figure S2 (related to Figure 4)**

Evaluation of the oxidative stress response of endogenous EIF6 cysteine in DU145 and HeLa cells. The cells were treated with 50 µM MND for 1 h. Black and white arrows show fully oxidized and reduced forms, respectively.

**Supplementary Figure S3 (related to Figure 5)**

Scatter plots showing quantitative data from the FLAG-IP MS analysis of RPL12 (A), MRPL50 (B), and TSFM (C). X-axis represents log_2_ fold change of intensities of proteins in FLAG-IP fraction prepared from HeLa cells transfected with bait-expressing plasmid compared to those transfected with empty vector (EV). Y-axis represents log_2_ fold change of intensities of proteins in FLAG-IP fraction prepared from bait-expressing cells treated with 50 µM MND for 1 h compared to those treated with DMSO. Proteins that were significantly more abundant in the bait FLAG-IP fraction than in EV are indicated by blue dots. The names of the baits and proteins reported to form the complexes are shown in the graph.

**Supplementary Figure S4 (related to Figure 5)**

Fluorescent images showing the localization of RPL12, MRPL50, TSFM, and MRPL50 in HeLa cells treated with DMSO, 50 µM MND or 50 µM MND plus 5 mM NAC. The cells were counterstained with Hoechst33342 to detect the nuclei. Scale bar = 10 µm.

## Supplementary Tables

**Supplementary Table S1**

All identified peptides by DIALRP in DMSO- and MND-treated DU145 cells

**Supplementary Table S2**

Quantitation data of 10,543 cysteine peptides

**Supplementary Table S3**

Oxidation percentage of 3,671 cysteine peptides

**Supplementary Table S4**

Statistical analysis data of 3,671 cysteine peptides using Perseus

**Supplementary Table S5**

Quantitation data of protein identified by DIALRP in DMSO- and MND-treated DU145 cells

**Supplementary Table S6**

DIVID functional annotation clustering of 295 highly oxidized proteins

**Supplementary Table S7**

List of 36 proteins annotated with GO translation.

**Supplementary Table S8**

Quantitation data of protein identified by FLAG-IP-MS analysis

**Supplementary Table S9**

Statistical data analysis of EIF2S2 FLAG-IP-MS using Perseus

**Supplementary Table S10**

Statistical data analysis of EIF6 FLAG-IP-MS using Perseus

**Supplementary Table S11**

Statistical data analysis of EFF2 FLAG-IP-MS using Perseus

**Supplementary Table S12**

Statistical data analysis of RPL12 FLAG-IP-MS using Perseus

**Supplementary Table S13**

Statistical data analysis of MRPL50 FLAG-IP-MS using Perseus

**Supplementary Table S14**

Statistical data analysis of TSFM FLAG-IP-MS using Perseus

**Supplementary Table S15**

Quantitation data of protein in MND-treated HeLa cells

