## Supplementary Figures for "Data-Independent Acquisition (DIA)-Based Label-Free Redox Proteomics (DIALRP) Reveals Novel Oxidative Stress Responsive Translation Factors"

**Supplementary Figure S1**

| | DMSO_rep1 | DMSO_rep2 | DMSO_rep3 | DMSO_rep4 |
|---|---|---|---|---|
| DMSO_rep1 | | 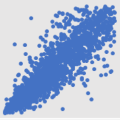 | 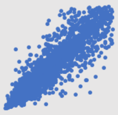 | 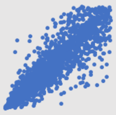 |
| DMSO_rep2 | 0.840 | | 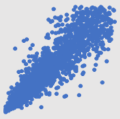 | 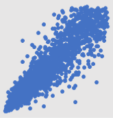 |
| DMSO_rep3 | 0.856 | 0.861 | | 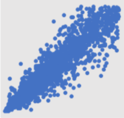 |
| DMSO_rep4 | 0.841 | 0.869 | 0.887 | |

### Supplementary Figure S2

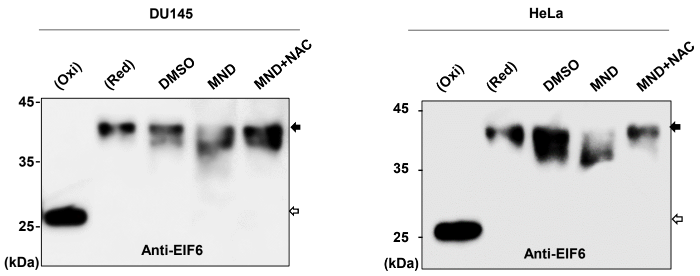

### Supplementary Figure S3

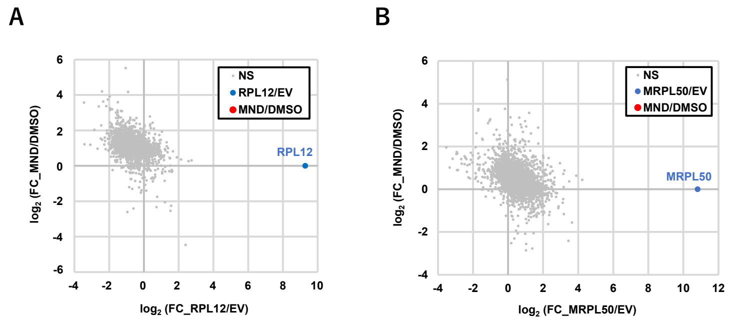

### Supplementary Figure S4

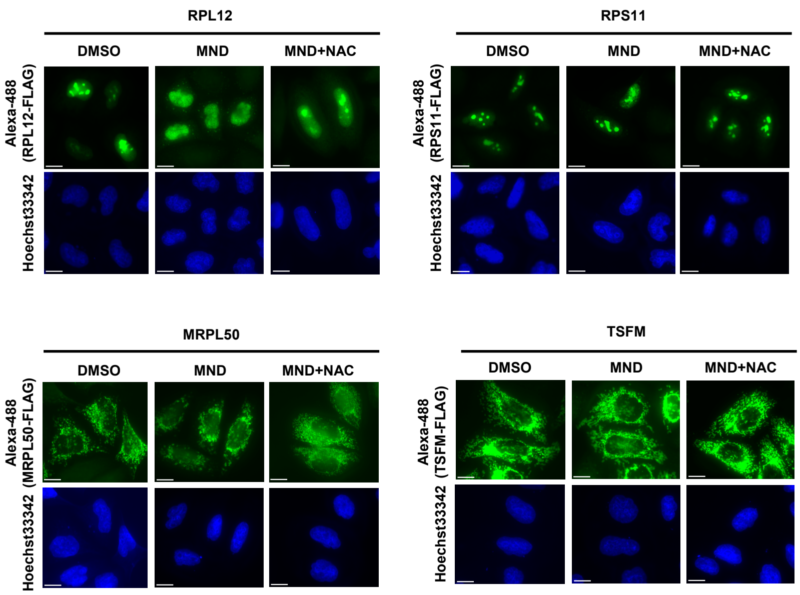
